# Extensive modulation of a conserved *cis*-regulatory code across 589 grass species

**DOI:** 10.1101/2025.04.23.650228

**Authors:** Charles O. Hale, Sheng-Kai Hsu, Jingjing Zhai, Aimee J. Schulz, Taylor Aubuchon-Elder, Germano Costa-Neto, Allen Gelfond, Mohamed El-Walid, Matthew Hufford, Elizabeth A. Kellogg, Thuy La, Alexandre P. Marand, Arun S. Seetharam, Armin Scheben, Michelle Stitzer, Travis Wrightsman, M. Cinta Romay, Edward S. Buckler

## Abstract

The growing availability of genomes from non-model organisms offers new opportunities to identify functional loci underlying trait variation through comparative genomics. While *cis*-regulatory regions drive much of phenotypic evolution, linking them to specific functions remains challenging. We identified 514 *cis*-regulatory motifs enriched in regulatory regions of five diverse grass species, with 73% consistently enriched across all, suggesting a deeply conserved regulatory code. We then quantified conservation of specific motif instances across 589 grass species, revealing widespread gain and loss over evolutionary time. Conservation declined rapidly over the first few million years of divergence, yet ∼50% of motif instances were conserved back to the origin of grasses ∼100 million years ago. Conservation patterns varied by gene class, with modestly higher conservation at transcription factor genes. To test for adaptive *cis*-regulatory changes, we used phylogenetic mixed models to identify motif gains and losses associated with ecological niche transitions. Our models revealed polygenic adaptation across 810 motif-orthogroup combinations, including convergent gains of HSF/GARP motifs at an Alpha-N-acetylglucosaminidase gene associated with adaptation to temperate environments. Our results support a “stable code, variable sites” model in which *cis*-regulatory evolution involves extensive turnover of individual binding site instances while largely preserving transcription factors’ binding preferences. *Cis*-regulatory changes at hundreds to thousands of genes appear to contribute to environmental adaptation. Our results highlight the potential of comparative genomics and phylogenetic mixed models to reveal the genetic basis of complex traits.

## INTRODUCTION

*Cis*-regulatory changes are arguably the most important genetic mechanism of evolutionary innovation, particularly over relatively shallow evolutionary time scales (King and Wilson 1975; Carroll 2008). While the evolution of *trans*-acting regulators such as transcription factors (TFs) typically have widespread effects genome-wide (Signor and Nuzhdin 2018), the evolution of *cis*-regulatory regions enables finely-tuned gene expression evolution at individual genes (Wittkopp et al. 2004; Prud’homme et al. 2007; Wray 2007). Variation within noncoding regions explains roughly half of additive trait variance within maize populations (Rodgers-Melnick et al. 2016), underscoring the importance of these regions in driving phenotypic evolution within species.

TF binding sites (TFBS) are key components of non-coding regulatory regions (Schmitz et al. 2021). TFBS are typically composed of a core DNA sequence motif that is recognized and bound by its cognate transcription factor. In plants, TFBS variants have been shown to be associated with the evolution of traits including inflorescence branching (Hendelman et al. 2021), abiotic stress tolerance (Jiang et al. 2022; Zeng et al. 2025) and fruit shape (Hu et al. 2024). Variants in the 5’ UTR and proximal promoter region are often particularly important for modulating expression levels (Cui et al. 2023; Voichek et al. 2024). Despite the importance of these TF binding variants in driving phenotypic evolution, characterizing key variants remains a major challenge. TF binding assays such as Chromatin Immunoprecipitation Sequencing (ChIP-seq), MNase-defined cistrome-Occupancy Analysis (MOA-seq), and DNA affinity purification sequencing (DAP-seq) have been very helpful for precisely characterizing where TFs bind throughout plant genomes (O’Malley et al. 2016; Savadel et al. 2021; Marand et al. 2022), but adapting them to work in non-model systems can be difficult. In particular, the increasing number of sequenced taxa requires scalable methods to discover key *cis*-regulatory features.

Characterizing evolutionary constraint and convergence across large numbers of taxa has emerged as a powerful strategy for determining genetic function from DNA sequence alone (Smith et al. 2020). The identification of conserved noncoding sequences has been used to identify highly important *cis*-regulatory features across non-model taxa (Haudry et al. 2013; Song et al. 2021; Stitzer et al. 2025). A limitation of this approach, however, is that many key *cis*-regulatory regions cannot be aligned reliably, even when function is shared (Schmidt et al. 2010; Yang et al. 2015; Kaplow et al. 2023). Alignment-free characterization approaches can offer a broader view of *cis*-regulatory evolution (Kaplow et al. 2023).

Due to these advances, our understanding of *cis*-regulatory evolution and its phenotypic implications is rapidly increasing. Still, some of the fundamental principles of regulatory evolution remain incompletely understood. A growing body of evidence indicates that TFs and their binding site preferences remain largely intact within lineages, suggesting a conserved “regulatory code” (Stergachis et al. 2014; Nitta et al. 2015; Tu et al. 2020). Changes to the regulatory code can occur via expansions or contractions of TF families that lead to the birth or extinction of particular TF binding preferences (Lehti-Shiu et al. 2017). Across deep evolutionary divergence, such as between plants and animals, widespread differences are observed between TFs and their binding preferences (Riechmann et al. 2000). The field lacks a clear consensus on the extent to which changes to TF binding preferences occur over varying time scales, as well as how and when changes occur. A deeper understanding of regulatory code evolution will inform how to effectively transfer genetics across species.

Another emerging model of regulatory evolution is that while TF binding preferences tend to be deeply conserved, individual instances of TF binding sites are extensively gained and lost over evolutionary time. We refer to this as the “stable code, variable sites” hypothesis.

Previous studies have estimated that only 20% of TFBS instances are conserved between humans and mice (Stergachis et al. 2014), and 20-40% of GOLDEN2-LIKE binding sites were found to be conserved between maize and rice (Tu et al. 2022). Relatively few studies so far have traced TFBS evolution across a large number of taxa (but see (Andrews et al. 2023)), or associated particular gain and loss events with diversification and adaptation at the macroevolutionary scale.

Grasses (Poaceae) have diversified widely over roughly 100 million years of evolution (Gallaher et al. 2022), occupying habitats ranging from tropical savannas to the Arctic tundra (Bouchenak-Khelladi et al. 2010; Edwards and Smith 2010; Spriggs et al. 2014; Lehmann et al. 2019). Grasses’ ecological diversification has been intertwined with repeated innovations in life history strategies (annual vs. perennial) (Kellogg 2015), photosynthetic pathway transitions (Grass Phylogeny Working Group II 2012), and photoperiod changes (Fjellheim et al. 2014).

Despite these significant shifts, aspects of genome structure such as gene content and collinearity are largely preserved across grass lineages (Bennetzen and Freeling 1993; McSteen and Kellogg 2022; Mascher et al. 2024), although ploidy (Stebbins 1985; Zhang et al. 2024), genome size (Bennett and Smith 1976) and gene regulatory patterns (Meng et al. 2021; Stitzer et al. 2025) vary widely across the family. Comparative genomic analyses across grasses are therefore well-suited to illuminate how broad and repeated environmental adaptation unfolds in complex genomes (Buell 2009).

We hypothesized that polygenic gain/loss of individual TFBS across the genome—rather than changes at a few master regulators—underlies grass diversification, enabling grasses to fine-tune gene regulation while maintaining regulatory logic. To test this hypothesis, we performed large-scale comparative genomic analyses across a collection of 727 genome assemblies representing 589 grass species. After establishing that grasses share a conserved set of TF motifs enriched in *cis*-regulatory regions, we quantified gain and loss of motif instances and performed association mapping across species to identify examples of motif gain and loss associated with environmental adaptation. We documented widespread gain and loss of motif instances as grasses adapted and diversified. Additionally, we revealed 810 environmentally associated motif gain/loss instances with moderate to weak levels of convergence. Together, our findings support a model of *cis*-regulatory evolution in which adaptation proceeds over macroevolutionary time scales via hundreds to thousands of *cis-*regulatory changes at small-effect downstream genes rather than at a few large-effect regulators.

## RESULTS

### Grasses share a deeply conserved *cis-*regulatory code

We hypothesized that a highly similar repertoire of *cis*-regulatory motifs are enriched in *cis*-regulatory regions across grasses. We estimated *cis*-regulatory regions using unmethylated regions (UMRs) available from five species (*Brachypodium distachyon, Oryza sativa*, *Sorghum bicolor*, *Zea mays*, and *Hordeum vulgare*). These five species last shared a common ancestor before the BOP (Bambusoideae, Oryzoideae, and Pooideae) and PACMAD (Panicoideae, Aristidoideae, Chloridoideae, Micrairoides, Arundinoideae, and Danthonioideae) clades diverged approximately 80 million years ago (Gallaher et al. 2022) (Fig. 1a). UMRs stably mark functional regulatory and genic regions of plant genomes and are rich in transcription factor binding sites (Crisp et al. 2020). To determine which *cis*-regulatory motifs are commonly enriched in regulatory regions, we quantified UMR enrichment of 704 experimentally-derived plant TF binding motifs from the JASPAR 2024 database (Rauluseviciute et al. 2023). Using randomized UMR sequences with preserved dinucleotide frequencies as background, we measured enrichment of each JASPAR motif in true UMRs relative to dinucleotide-shuffled UMRs.

**Figure 1.**
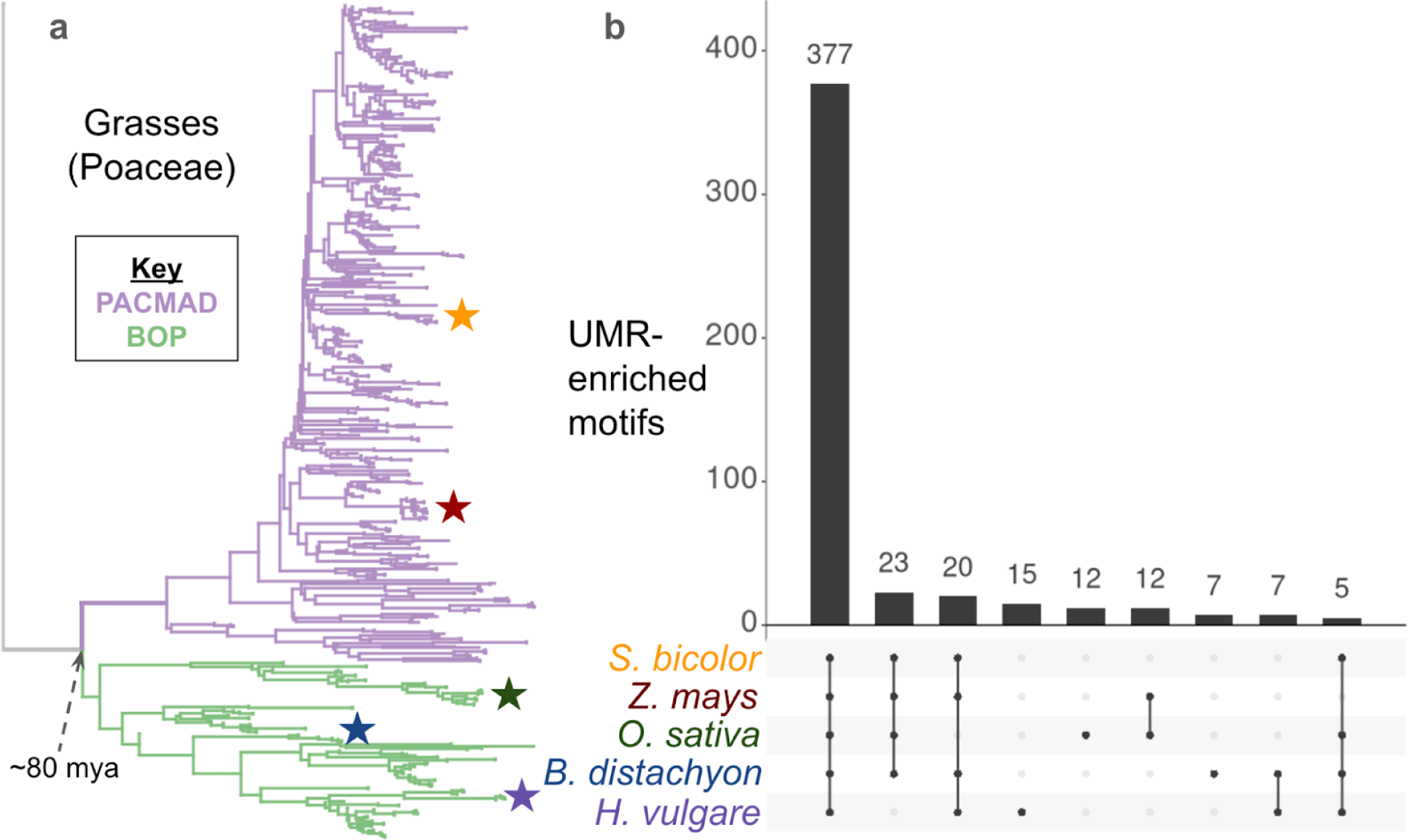
Grasses share a conserved set of UMR-enriched motifs. (a) Phylogenetic tree of 589 Poaceae species. Five representative species are starred: *Sorghum bicolor* (orange), *Zea mays* (red), *Oryza sativa* (green), *Brachypodium distachyon* (blue), *Hordeum vulgare* (purple). (b) Enrichment of transcription factor motifs in unmethylated regions across five representative grass species. The intersection bars show the number of enriched motifs for each species set. Intersections with fewer than five motifs are not shown.

Enrichment fold-change showed correlations of 96-99% between *B. distachyon, O. sativa,* and *S. bicolor*, as well as with *Zea mays* and *Hordeum vulgare* (Fig. S1). Of the 514 motifs that were enriched in at least one species, 73% (377 / 514) were commonly enriched across all five species (Fig. 1), with a similar set of motifs enriched in accessible chromatin regions across species (Fig. S2). We used this set of 377 commonly enriched motifs (Supplementary Table 1) for subsequent comparisons of motif conservation and turnover across the grass family.

### Patterns of motif occurrence across 589 widely-adapted species

To characterize *cis*-regulatory evolution on a large scale, we built a pipeline to quantify and compare motif occurrences across orthologous regions of hundreds of species (Fig. 2a). For this study, we used publicly-available WGS short reads to generate genome assemblies for 57 species. While not highly contiguous, our WGS assemblies recovered a median of 4206 genes (75%) from a set of 5592 conserved grass genes in a BUSCO-like analysis (Fig. S3b, Supplementary Table 2). In total, we amassed a dataset of 727 genome assemblies representing 589 diverse grass species using the 57 new WGS assemblies along with 211 public genome assemblies, 368 newly generated short read assemblies from Schulz et al (unpublished data), 58 short read assemblies from (Schulz et al. 2023) and 33 highly contiguous assemblies from (Stitzer et al. 2025) (Fig. S3a, Supplementary Table 2). Our dataset captures wide and repeated environmental adaptation across the grass family (Hsu et al, unpublished data).

**Figure 2.**
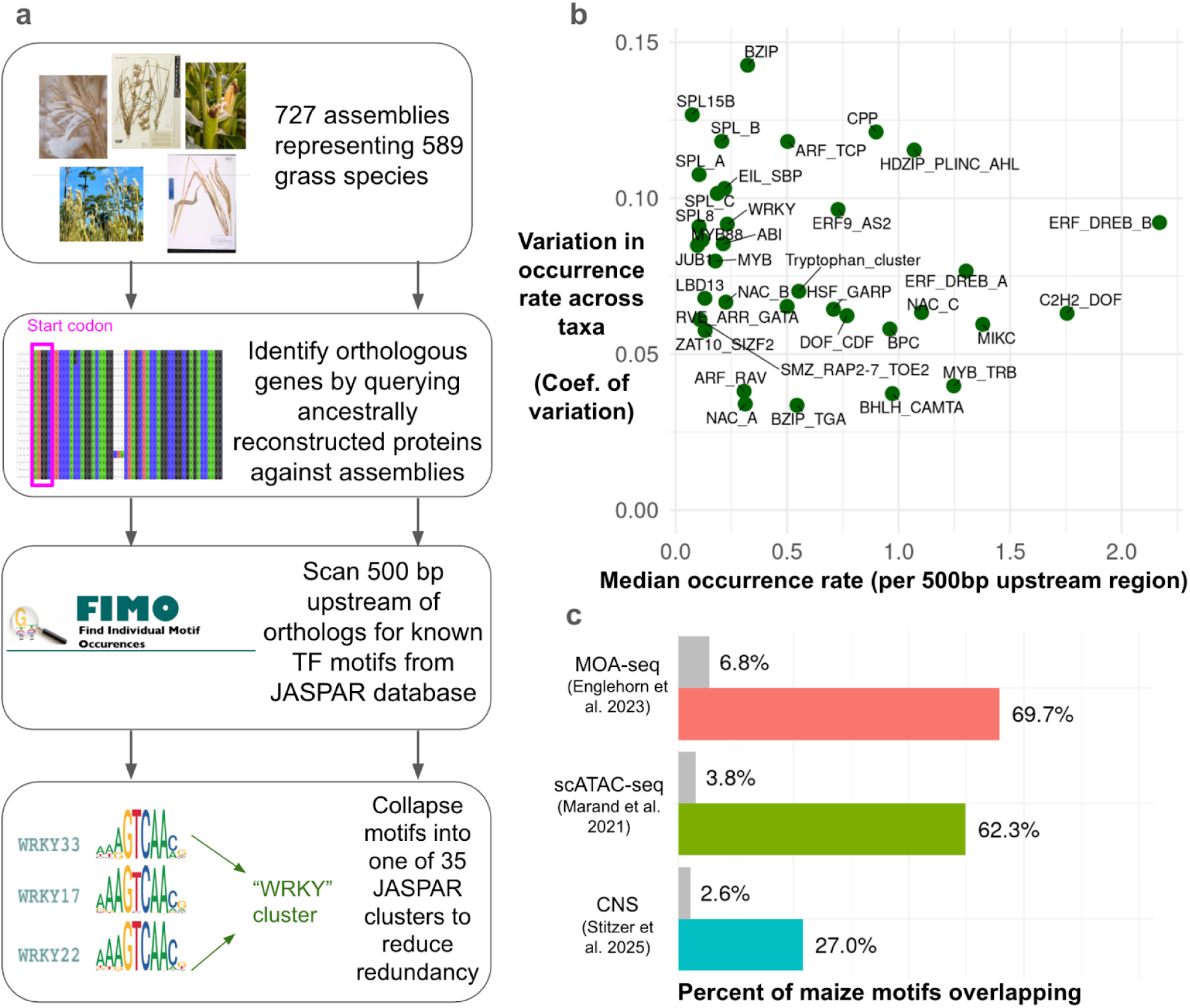
Summary of TF motif profiling across 727 assemblies. (a) Depiction of bioinformatic pipeline to identify TF motif instances upstream of orthologous genes across taxa. (b) Occurrence rate and variation across species for each motif cluster (c) Overlap of maize upstream motifs with known *cis*-regulatory features. Colored bars show the percentage of motifs overlapping intervals from each feature type, while grey bars represent the mean overlap percentage with randomly shuffled intervals.

To enable cross-species comparisons, we identified orthologous genes across taxa by querying a set of ancestrally reconstructed protein sequences against each assembly and retaining the primary alignment. We retained 15,555 orthogroups after filtering, each containing an average of 465 taxa (SD = 104) (Fig. S4), indicating a high level of taxonomic representation suitable for robust comparative analyses. We scanned intervals 500 bp upstream of the aligned translation start sites for the set of 377 motifs with conserved UMR enrichment. While many important regulatory elements are present further upstream (or downstream) from genes, we considered only 500 bp upstream due to 1) limited contig length in our short-read assemblies and 2) the higher proportion of transposable elements present further upstream. We collapsed overlapping instances of similar motifs into a single merged interval to reduce redundancy for subsequent analyses, using 35 motif clusters delineated by JASPAR2024 (Supplementary Table 1).

The 35 motif cluster types differed widely in abundance and variability across taxa (Fig 2b). Most motif clusters had a median occurrence rate of ∼0.25 instances per upstream region, but some motif clusters such as those containing ethylene-responsive factors (ERF) and dehydration responsive element binding proteins (DREB) were highly abundant, appearing upwards of twice per upstream region on average. Occurrence rate variability across species also differed widely across motif clusters, with coefficients of variation ranging from 0.03 to 0.14 (Fig 2b).

One challenge of *de novo* TFBS prediction using motifs is that many false positives are typically observed when scanning genome-wide (Grant et al. 2011; Jayaram et al. 2016).

Restricting motif scanning to regions with external functional evidence (e.g. regions with evolutionary conservation, open chromatin, or in this case, gene proximity) can reduce the number of false positives (Wasserman and Sandelin 2004), though motifs remain unreliable predictors of *in vivo* binding (Krieger et al. 2022; Mahendrawada et al. 2025). To investigate overlap of our set of gene-proximal motifs with known regulatory regions, we quantified the proportion of our maize motif instances found within independently generated maize MOA-seq (TF binding) peaks (Engelhorn et al. 2023), accessible chromatin regions from a single-cell assay for transposase accessible chromatin with sequencing (scATAC-seq) (Marand et al. 2021), and conserved noncoding sequences (CNS) (Stitzer et al. 2025). The majority of maize motifs intersected scATAC-seq (62.1%) and MOA-seq peaks (69.8%), while only 26.8% intersected CNS (Fig. 2c). While our focus on proximal upstreams prevented us from examining the multitude of distal TF binding sites that can be captured by CNS and ‘omics approaches, our approach captures a set of motifs in active regulatory regions that are not alignable and may be missed by CNS analyses. For example, we were able to recover a known functional MYB TFBS at *ZmICE1 (Jiang et al. 2022)* that is not found within a CNS (Fig. S5).

### Motif conservation decays nonlinearly with increasing evolutionary divergence

To track the evolutionary gain and loss of motif instances in grasses, we measured motif conservation by comparing the number of motifs found at maize orthologs with those found in orthologs of the other 588 species (see Methods). The decay in motif conservation relative to maize was extensive. Roughly 60% of maize motif instances were shared with *Sorghum* (∼15 million years diverged) at orthologous upstream regions, and 50% were shared with rice across ∼80 million years of evolution (Fig. 3a). On average, motif conservation between orthologs did not decay to the level observed between random maize genes, approaching an asymptote of 48% conservation (Fig. 3a). Decay curves were similar when defining motif conservation relative to *Sorghum* (Fig. S6a) and rice (Fig. S6b) instead of maize. Additionally, motifs that co-localized with maize ChIP-seq peaks showed minimal evidence of elevated conservation (Fig. S7), indicating that our motif conservation estimates were not substantially biased by the large number of motifs lacking *in vivo* binding support.

**Figure 3.**
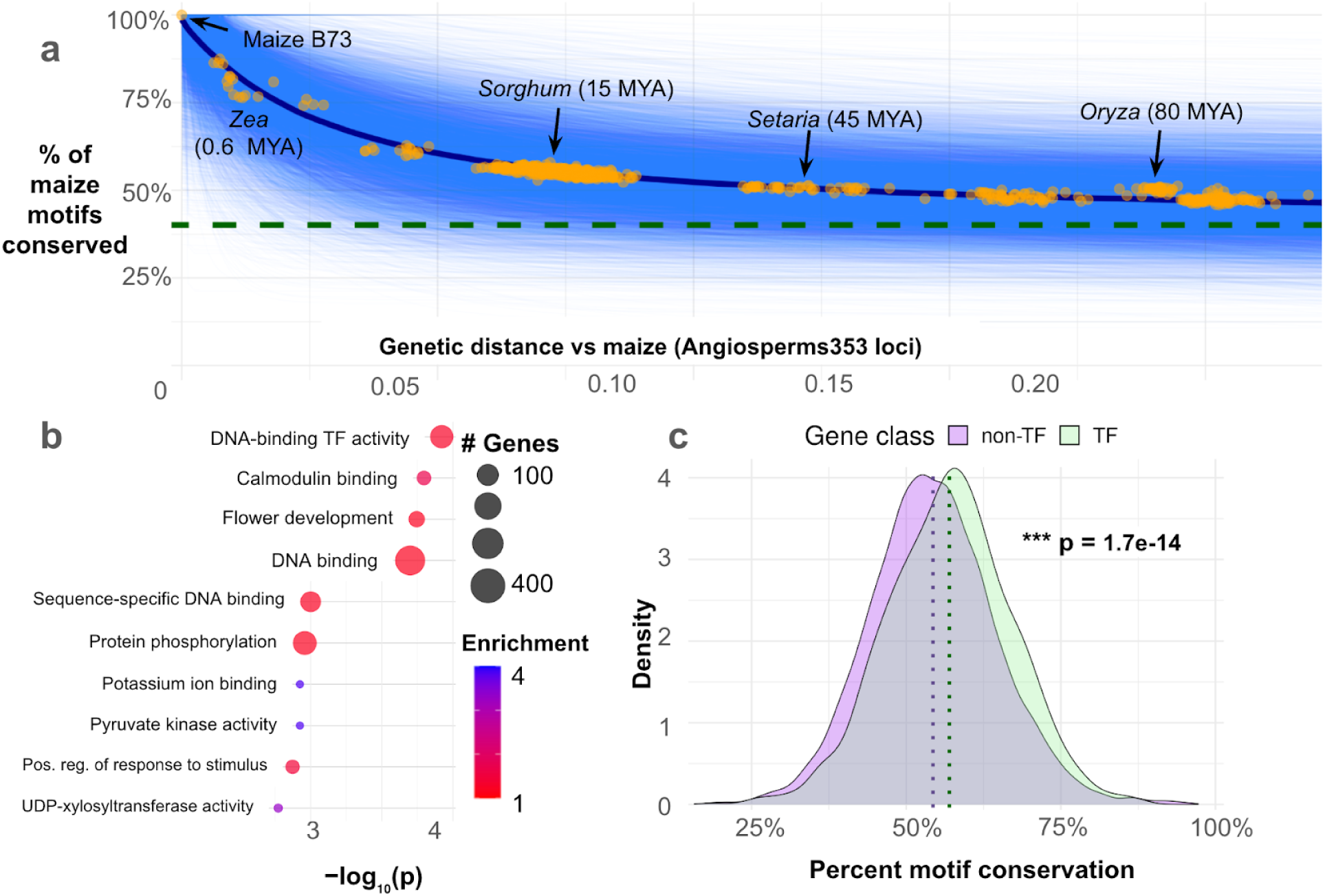
Motif conservation decays nonlinearly and heterogeneously across gene classes. **a.** Percentage of maize motif instances conserved across 589 Poaceae species. Genetic distance was estimated using pairwise distances between maize and Poaceae species at the Angiosperms353 loci. Blue lines represent exponential decay curves fit for 12,784 orthogroups. Orange points show the mean percentage of maize motifs conserved in each Poaceae species across all orthogroups, with an exponential decay curve depicted in dark blue. Dashed green line represents the mean percentage of motifs conserved across 100,000 random pairs of maize genes. Approximate divergence times from maize are shown for key taxa (Chen et al. 2022; Gallaher et al. 2022)**. b.** Enriched gene ontology terms for the orthogroups in the top quartile of motif conservation. **c.** Motif conservation at transcription factor orthogroups (n = 1,204) versus at non-transcription factor orthogroups (n = 11,580). Median values for each class are shown by dotted vertical lines. P-value is from an asymptotic two-sample Kolmogorov-Smirnov test.

Conservation of motif instances varied widely by gene, with most orthogroups approaching 40-60% conservation between the most distantly related grasses (Fig 3a). Coding sequence conservation explained minimal variance in motif conservation at individual orthogroups (R-squared = 0.03) (Fig. S8). A small number of highly conserved proteins, such as ribosomal and plastid-encoded proteins, exhibited very high rates of motif conservation.

However, the motifs detected near plastid genes are likely artifacts of motif scanning, as plastid genes are not regulated by nuclear transcription factors. Orthogroups with higher motif conservation were significantly enriched (Fisher’s exact test, p < 0.05) for 121 gene ontology (GO) terms. Many of the strongest enrichments were related to regulatory function by transcription factors (Fig. 3b, Supplementary Table 3). Specifically, TF genes (n = 1201) exhibited slightly higher motif conservation than non-TFs (n = 11,537) (TF median = 57%, non-TF median = 54%; Kolmogorov-Smirnov test, p = 1.7e-14) (Fig. 3c). The difference between TFs and non-TFs persisted when comparing motif conservation between maize and sorghum at syntenic orthologs only (Fig. S9).

In addition to regulatory functions, we observed many significant GO terms among high-conservation genes related to signal transduction (e.g. “calmodulin binding”, “protein phosphorylation”, and “phosphorelay signal transduction system”), cytoskeleton (e.g. “actin filament organization”, “microtubule cytoskeleton organization”, and “cortical microtubule organization”), and development (“flower development”, “abaxial cell fate specification”, and “positive regulation of long-day photoperiodism, flowering”). Low-conservation genes were enriched for 110 terms including “defense response”, “plant-type cell wall”, and “RNA modification” (Fig. S10, Supplementary Table 3). GO terms that were not significantly enriched among either high or low-conservation genes included “response to abiotic stimulus”, “photosynthesis”, and “reproduction”.

### Convergent motif gain/loss occurs at hundreds of genes during environmental adaptation

To investigate which motif instances are linked to environmental adaptation, we employed phylogenetic mixed models, which can be used to control for phylogenetic relatedness while associating genomic features with traits across species (Housworth et al. 2004). We characterized each species’ ecological niche as described in (Hsu et al. 2024), summarizing into ten environmental principal coordinate (envPC) axes that together explain 75% of total ecological variability among the natural environments of grasses across the globe.

We hypothesized that repeated gains or losses of particular motifs underlie environmental adaptation in grasses. We first tested whether global occurrence rates of motifs across taxa are associated with environmental adaptation, using the envPCs as proxies of environmental diversity (Fig. 4a). Since variable genome sizes have been hypothesized to alter the spacing of regulatory regions (Schmitz et al. 2021), we controlled for the overall density of all motifs as well as the background mono and di-nucleotide content in genomic background regions, which could influence the probability of detecting a motif by chance. Occurrence rates of C2H2-type zinc finger (C2H2/DOF), DNA binding with one finger (DOF/CDF), NAM, ATAF and CUC (NAC), and homodomain-leucine zipper / Plant Zinc Finger / AT-HOOK MOTIF NUCLEAR LOCALIZED (HDZIP/PLINC/AHL) motifs were significantly associated with envPC1 (FDR-adjusted Wald test, p < 0.01) (Fig. 4b, Supplementary Table 4), which primarily captures thermal variability across taxa. Additionally, the occurrence rate of basic leucine zipper (BZIP) motifs was significantly associated with envPC2, which captures variation in moisture availability.

**Figure 4.**
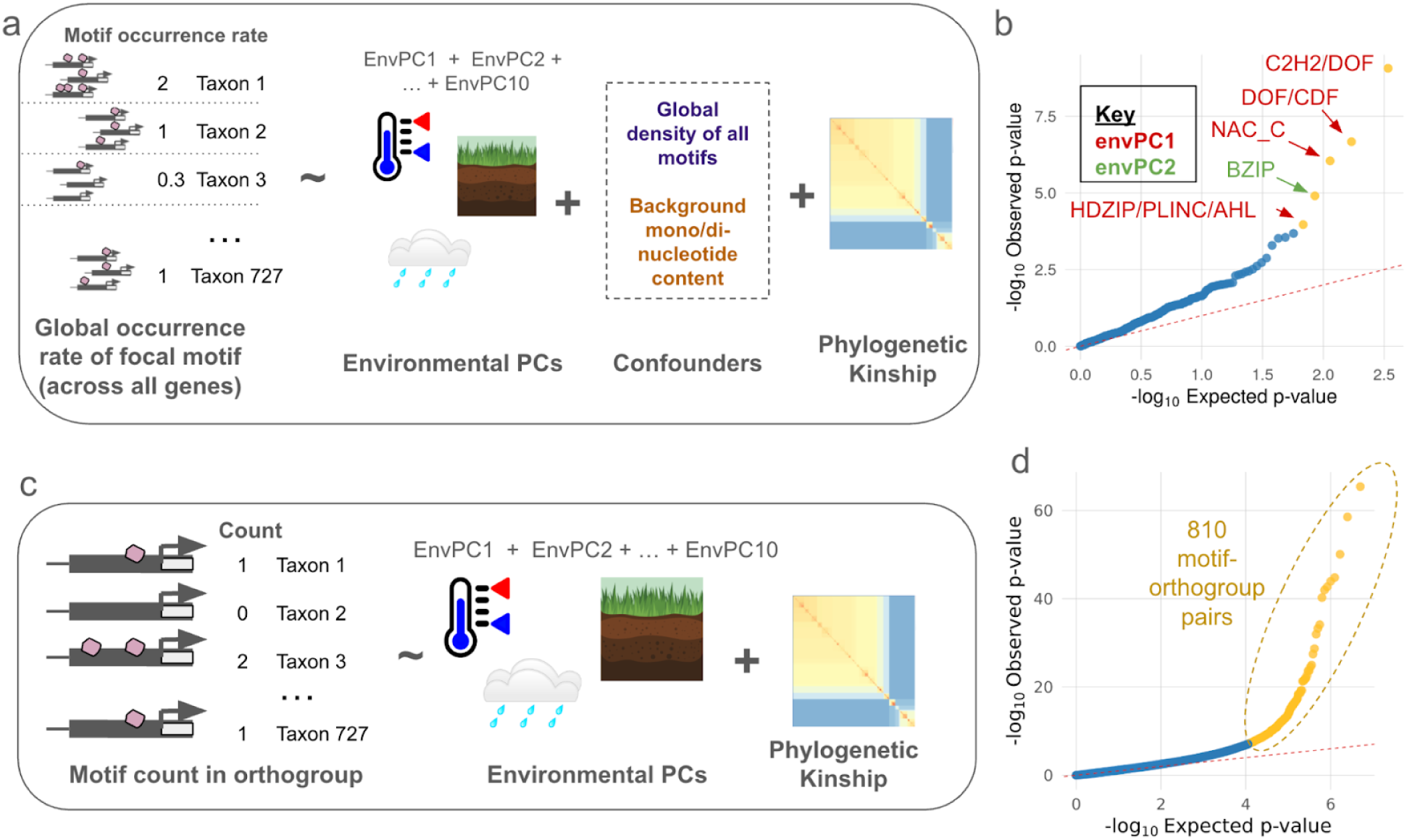
Polygenic turnover at hundreds of motif instances is associated with environmental adaptation.**a.** Setup of global occurrence association models. One phylogenetic mixed model was run for each of the 35 motif types. **b.** Quantile-quantile plot showing observed p-values from the global occurrence models versus p-values expected under the null hypothesis. Observed p-values were calculated from Wald tests across 350 fixed effect envPC terms. Terms with FDR-corrected p-values < 0.01 are shown in yellow. **c.** Setup of orthogroup-specific models. One model was run for each motif-orthogroup combination for a total of 498,287 models. **d.** Quantile-quantile plot for orthogroup-specific models. Observed p-values were calculated for ∼5 million fixed effect envPC terms.

We further hypothesized that repeated gains or losses of motif instances at particular orthogroups are associated with environmental adaptation. For each motif-orthogroup combination we associated the number of motifs per assembly at an orthogroup with the ten environmental PCs after controlling for phylogenetic relatedness (Fig. 4c). Across 498,287 models, we scanned for signals of convergent motif gain or loss. We identified 810 unique motif-orthogroup combinations (1025 total) with strongly significant (FDR-adjusted Wald test, p < 0.01) environmental associations (Fig. 4d, , Supplementary Table 5), suggesting convergent motif gain or loss across independent lineages.

### Gain of HSF/GARP motifs at an Alpha-N-acetylglucosaminidase gene predicts cold adaptation

We reasoned that the orthogroups most strongly associated with envPCs would be enriched for pathways and processes linked to abiotic stress responses. Indeed, a GO enrichment analysis of the 810 significant orthogroups identified 47 enriched terms, many of which were related to known abiotic stress response processes such as oxidative stress response (“cellular oxidant detoxification”, “response to oxidative stress”, “oxidoreductase complex”, “oxidoreductase activity”) and degradation of misfolded proteins (“ubiquitin-dependent ERAD pathway”, “proteolysis involved in protein catabolic process”, “ubiquitin-specific protease binding”, “endoplasmic reticulum unfolded protein response”, “threonine-type endopeptidase activity”) (Fig. 5a, Supplementary Table 6).

**Figure 5.**
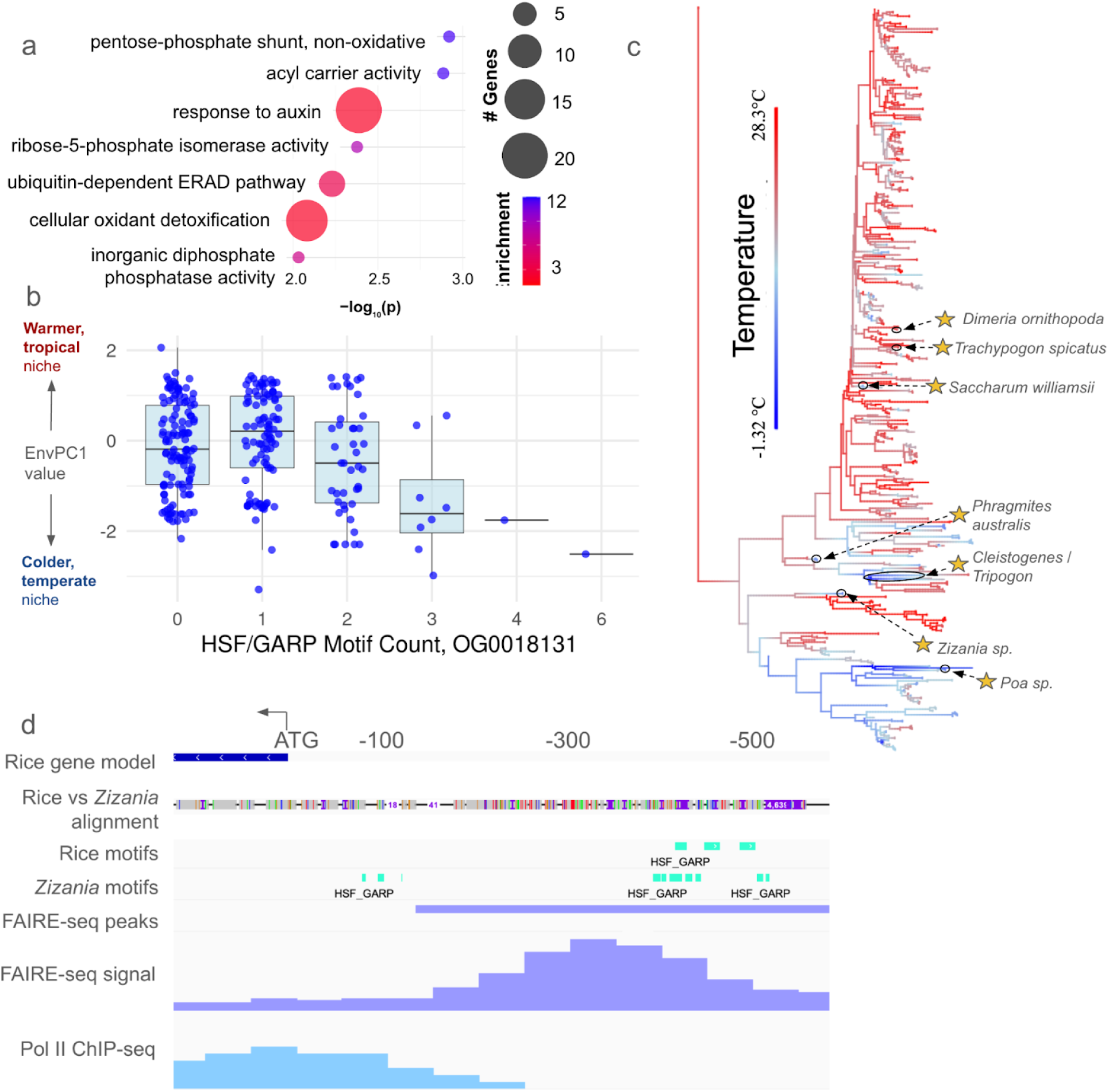
Exploration of top motif-orthogroup pairs. **a.** GO enrichment analysis for 810 significantly-associated orthogroups. **b.** Environmental PC1 values for taxa with varying numbers of HSF/GARP motifs at OG0018131. **c.** Repeated gain and loss of HSF/GARP motifs at OG0018131 across the grass family. Taxa with three or more copies of HSF/GARP are starred. **d.** HSF/GARP motif variation between *Z. palustris* and *O. sativa* at an open chromatin region upstream of OG0018131.

We investigated some of the individual motif-orthogroup combinations related to these enriched processes. One of the motif-orthogroup combinations most strongly associated with envPC1 was a Heat Shock Factor / Golden2, ARR-B, Psr1 (HSF/GARP) motif at OG0018131 encoding an Alpha-N-acetylglucosaminidase family protein. Alpha-N-acetylglucosaminidase is involved in N-glycan degradation (Ronceret et al. 2008), which has been connected to protein misfolding response under chilling in Arabidopsis (Ma et al. 2016) and alfalfa (Xu et al. 2024)). Additionally, HSF TFs are strongly implicated in thermal stress responses (Guo et al. 2016), and in grasses various HSFs have been linked to misfolded protein responses (Cheng et al. 2015) and chilling tolerance (Gao et al. 2024).

Increasing counts of HSF/GARP motifs at OG0018131 predicted lower envPC1 values (Fig. 5b), with the association primarily driven by taxa containing three or more HSF/GARP instances. At least seven independent lineages had three or more copies of the motif (Fig. 5c), representing frost-tolerant genera including *Zizania* (wild rice), *Phragmites*, and *Poa*. Notably, only a fraction of independent cold adaptation events (very roughly 25%) were associated with expansions of this motif, indicating that the strength of convergence at this gene was relatively weak despite strong statistical support (Wald test, p =1.89e-10). Rice contains multiple copies of the motif in an open chromatin region, and its cold-tolerant relative *Zizania palustris* contains additional copies (Fig. 5d).

## DISCUSSION

Our results support a “stable code, variable sites” model of *cis-*regulatory evolution under which transcription factor binding preferences remain conserved across deep divergence while individual TF binding sites across the genome are frequently gained and lost. The high overlap of UMR-enriched motifs across species suggests that minimal lineage-specific loss of binding preferences has occurred in grasses, consistent with the deep conservation of DNA binding domains observed within many transcription factor families (Yamasaki et al. 2008; Weirauch and Hughes 2011; de Mendoza et al. 2013). Importantly, our analysis does not rule out lineage-specific gains of binding site preference (perhaps via diversification of transcription factor families), which could be investigated using *de novo* motif characterization. Quantitative estimates of binding affinity would be needed to more sensitively detect subtle shifts in binding preferences. Regardless, the high conservation of motif sequences we observed across distantly related species highlights that TFs in grasses share similar binding preferences, likely with transferable regulatory functions across species. Our implicit assumption is that UMR enrichment is evidence of motif function, although motif depletion can also be meaningful since selection on *cis*-regulatory regions may act to inhibit particular TF binding events (He et al. 2011).

While much of grasses’ regulatory code appears to have been conserved across a hundred million years of evolution, our work demonstrates that extensive remodeling of individual *cis*-regulatory regions has occurred. Although some TF binding site instances have been shown to be as strongly conserved as coding sequences (Heyndrickx et al. 2014), just 50% of the motif instances we observed were conserved between maize and rice, which are roughly as genetically divergent as humans and mice (Gallaher et al. 2022). Despite extensive turnover overall, conservation of some motif instances appears to persist over deep evolutionary time scales. Motif conservation at orthologous regions decays nonlinearly and tends to approach a minimum level of conservation between 40-60% for most grass orthogroups. This suggests that regulatory regions may often contain a handful of strongly-conserved motif instances alongside many motifs that are rapidly turned over.

Our finding that regulatory genes, and specifically transcription factors, have slightly higher motif conservation suggests that a large amount of cis-regulatory evolution within regulatory networks occurs at terminal target genes rather than just at a handful of key regulators. This finding supports a model in which transcription factors and other highly pleiotropic genes exhibit greater constraint, in line with past theoretical and empirical work (Prud’homme et al. 2007; Chesmore et al. 2016; Nocchi et al. 2024), but see (Khaipho-Burch et al. 2023). Future studies could explicitly test whether the degree of pleiotropy of a gene predicts conservation of its TF binding sites. Alternatively, background selection on the target gene could be driving conservation of proximal TFBS rather than selection on the TFBS itself, though the low correspondence between coding sequence and motif conservation suggests that this effect is minimal.

Our environmental association models suggest that *cis*-regulatory changes in grasses are not concentrated at a handful of key adaptation genes, but occur pervasively across a wide set of diversely functioning genes. We observed weakly convergent motif gain and loss at hundreds of genes with diverse molecular functions, underscoring that environmental adaptation involves altering regulation of a highly diverse set of genes, not just modifying a handful of pathways or processes. These findings are consistent with theoretical findings that suggest that global adaptation of complex traits tends to be associated with many small-effect regulatory changes (Prud’homme et al. 2007). This challenges attempts to leverage cross-species insights for crop genetic engineering. Still, characterization and editing of key TF binding sites remains a promising approach for precisely tuning expression levels.

Our results suggest that the phylogenetic mixed model may be an effective approach to nominate key candidate loci using large-scale comparative genomics. With more sequenced genomes available, the ability of such comparative genomic approaches to detect convergently evolving genome features will improve (Smith et al. 2020), particularly for loci of small effect. Future studies could build on this framework by pairing large-scale comparative genomics screens with detailed molecular characterization of the top candidate loci. For example, a variety of mechanisms could underlie the HSF/GARP motif variation we observed at OG0018131.

Gains of HSF/GARP motifs at this locus could alter TF binding affinity of HSF/GARP TFs to the region, or perhaps permit combinatorial binding of multiple distinct HSF/GARP TFs that recognize similar motifs. Further work could validate the functional effect of this motif and its relationship with environmental adaptation.

A number of tradeoffs were required to scale up our analyses across hundreds of complex plant genomes. The relatively high overlap of gene-proximal motifs intersecting open chromatin and TF binding regions suggests that many gene-proximal motifs are likely to be bound by transcription factors *in vivo*. Still, a significant fraction of the motif instances we characterized are likely to be non-functional. Additionally, studies have found that only a fraction of TF binding changes are due to motif variants (Reddy et al. 2012; Krieger et al. 2022), underscoring that TF binding assays, though far less scalable, currently offer more reliable estimates of TFBS than sequence-based prediction.

The low overlap we observed with conserved non-coding regions supports previous findings that alignment-based approaches, while useful for identifying high-importance regions from DNA sequence alone, can miss functional regulatory features (Yang et al. 2015; Kaplow et al. 2023). Alignment-free approaches such as deep learning models hold promise going forward for scalable characterization of *cis*-regulatory regions. The ability of such models to represent the orientation, positioning and co-binding context of *cis*-regulatory elements can offer a more nuanced picture of *cis*-regulatory evolution beyond simple gain and loss of motifs.

Our study focused on characterizing motif variants in proximal upstream regions due to limited assembly contiguity. However, changes to distal *cis*-regulatory elements (Clark et al. 2006) or *trans*-acting factors also contribute strongly to regulatory adaptation and were not considered in our analyses of motif gain and loss. Another limitation of our approach is that many duplicated gene copies, particularly in the numerous polyploid taxa represented in the dataset, were filtered out. Additionally, because we relied on orthogroups across wide phylogenetic distances to make comparisons, lineage-specific genes that may be key to adaptation were not considered. Similarly, the reliance on convergent evolution in our association models approach may miss adaptive mechanisms that are influenced by evolutionary contingencies and constraints; e.g. lineages using C4 photosynthesis may have different physiological strategies “available” to them than C3 lineages. Future comparative genomic analyses will benefit from complementing broad phylogenetic comparisons with comparisons of closely related species that largely share the same physiology and genetics. Moreover, integrating more detailed estimates of ecological context into population genetic analyses of key taxa would offer greater resolution and nuance beyond our broad species-level estimates of adaptation.

## CONCLUSION

We performed large-scale comparative genomic analyses across 589 grass species to investigate the *cis*-regulatory basis of diversification and environmental adaptation in grasses. We found that grasses share a deeply conserved regulatory code, with 377 *cis-*regulatory sequence motifs conserved across diverse species. We documented extensive gain and loss of specific motif instances across the grass family, suggesting extensive reorganization of orthologous

*cis*-regulatory regions. *Cis*-regulatory changes do not appear to be highly concentrated at particular classes of genes, though they occur slightly less frequently at regulatory genes such as transcription factors. 810 motif-orthogroup combinations show evidence of convergent gain and loss of motifs associated with environmental adaptation. However, even the most strongly significant motifs are only weakly repeatable, underscoring the diversity of *cis*-regulatory routes to environmental adaptation.

## MATERIALS & METHODS

*Transcription factor motif scanning:* We downloaded 805 TF motifs and clusters from the 2024 JASPAR CORE non-redundant plant collection (Rauluseviciute et al. 2023). This set of motifs has strong experimental evidence from TF binding assays such as DAP-seq and ChIP-seq primarily performed in *Arabidopsis*. All motif scans were performed using FIMO from the MEME suite v5.5.7 (Grant et al. 2011) with the parameters --max-strand --no-qvalue --skip-matched-sequence --max-stored-scores 100000000. Homogenous background mono- and di-nucleotide frequencies (=0.25 per nucleotide) were used to avoid species-specific biases in motif detection thresholds. 704 of the 805 motifs in the JASPAR collection were detectable using our parameters, with the rest containing insufficient information content to enable statistically significant matches with FIMO.

*UMR enrichment analysis:* Unmethylated regions (UMRs) for five diverse grass species (*Sorghum bicolor, Oryza sativa, Brachypodium distachyon, Zea mays, Hordeum vulgare*) were downloaded from (Crisp et al. 2020). Background sequences were generated by di-nucleotide shuffling each UMR region 100 times, preserving local sequence composition. Motif scanning was performed using FIMO on both UMR and background sequences. The fitdist function in R package “fitdistrplus” (Delignette-Muller and Dutang 2015) was used to fit probability distributions to background motif counts. Two-tailed p-values were then calculated to identify significantly over- and under-represented motifs in UMR regions. We quantified how many motifs were significantly over-represented (FDR-adjusted p value < 0.01 from Fisher’s exact test) within and between species. The 377 commonly-enriched motifs were used for subsequent analyses.

For accessible chromatin enrichment, we downloaded bulk accessible chromatin regions for *S. bicolor, O. sativa, B. distachyon, Z. mays, H. vulgare, Setaria viridis*, and *Arabidopsis thaliana* from (Lu et al. 2019) and performed enrichments as described above.

*Compiling genomic dataset:* We compiled a dataset of 727 genome assemblies representing 589 diverse grass species by combining 217 publicly-available assemblies from NCBI (Sayers et al. 2024), Phytozome (Goodstein et al. 2012), and CoGE (Lyons and Freeling 2008) with 33 long-read assemblies from (Stitzer et al. 2025) and 550 additional assemblies from short reads using the high-throughput assembly pipeline described in (Schulz et al. 2023) (Supplementary Table 2). For this study, we assembled 57 new short-read assemblies from WGS raw reads deposited in the NCBI SRA database (Sayers et al. 2024). We downloaded WGS data from all Poaceae species lacking an existing genome assembly if at least 15GB of WGS data was available for that species. We then assembled the SRA genomes *de novo* using Megahit v1.2.9 (Li et al. 2015) with minimum kmer size = 31 and default parameters for other assemblies, as described in (Schulz et al. 2023). If < 30GB WGS data was available for the largest WGS accession for a species, we merged WGS data from multiple accessions to improve assembly completeness. We performed three QC steps on our SRA assemblies. First, we ran Kraken2 v2.1.3 (Lu et al. 2022) on raw reads (subsampled to a depth of 10M reads) using the PlusPFP database to verify sample identify on a rough taxonomic scale and flag accessions with high levels of bacterial or fungal contamination. Second, we visualized the assemblies on a *matK* phylogenetic tree against labeled *matK* sequences from the BOLD database (Ratnasingham et al. 2024) to rule out obviously-mislabeled accessions. To generate *matK* alignments, we downloaded a complete *matK* CDS sequence for *Streptochaeta angustifolia* (Genbank: AF164382.1) (Hilu et al. 1999). We queried the *S. angustifolia* sequence against all WGS short read assemblies using minimap2 v2.17 (Li 2018) with the parameters -ax asm20 –eqx -I 100g –secondary=no, extracting the primary alignment in each assembly. Using the extracted *matK* sequences from our WGS assemblies and 7,338 Poaceae *matK* sequences from the BOLD database, we performed multiple alignment with mafft v7.520 (Katoh et al. 2002) --auto. Then we constructed a phylogenetic tree using raxml v8.2.13 (Stamatakis 2014) with the GTRGAMMA model. As a final assessment of assembly quality, we quantified assessed assembly completeness using TABASCO, a BUSCO-like assembly metric designed specifically for use with grasses (Schulz et al. 2023) which labels a set of 5592 query genes as “complete”, “duplicated”, “fragmented”, or “missing”.

*Orthogroup construction:* Across the 727 genome assemblies, we selected 32 representative high quality long read assemblies to construct orthogroups. In order to avoid potential annotation biases, we ran Helixer (Stiehler et al. 2021) to annotate each of the representative genomes and extracted the protein sequences. Based on protein sequence homology, orthogroups were constructed using OrthoFinder v2.6.4 (Emms and Kelly 2019). In total, we obtained 22,503 orthogroups with homologous sequences represented by more than eight of the representative genomes. From the multiple sequence alignment of each orthgroup, we reconstructed the ancestral protein sequence using the R/phangorn package (Schliep 2011). The ancestral sequences of the orthogroups were used to query the orthologs in all 727 genomes by miniProt v0.13.0 (Li 2023).

*Phylogenetic characterization:* We calculated phylogenetic relatedness among the 727 studied genomes using the Angiosperms353 loci (McDonnell et al. 2021). We identified the orthogroups homologous to the Angiosperms353 loci using miniProt (v0.13.0) and generated gene trees for each of the orthogroups using RAxML v8.12 (GAMMA + GTR). ASTRAL-Pro v2 (Zhang and Mirarab 2022) was used to reconcile the species tree based on the gene trees. We generated a matrix of shared branch length among any pair of tips in the species tree to represent the relatedness between them. This matrix (hereafter phyloK matrix) is included in our phylogenetic mixed model to account for shared macro-evolutionary history among species. Using the concatenated alignment of the Angiosperms353 loci, we calculated pairwise genetic distance among the 727 taxa based on the K81 model implemented in R/ape::dist.dna() function. We obtained divergence time estimates for *Zea-Tripsacum* from (Chen et al. 2022), and estimated divergence time across Poaceae from (Gallaher et al. 2022).

*Motif annotation in orthologous upstream regions:* Alignments were filtered to retain primary alignments that did not contain a frameshift or a premature stop codon, and that began with a start codon. To approximate 5’UTR and promoter regions across assemblies, 500 bp sequence was extracted upstream of the aligned start codon. 500 bp was chosen as a conservative estimate of 5’ UTR and promoter regions to maximize species representation in the dataset given the limited contig lengths for the short-read assemblies. If 500 bp upstream sequence was not available, or if the sequence contained > 5% Ns, the sequence was discarded from analysis.

Orthogroups with fewer than 200 assemblies represented were dropped. To minimize redundant, overlapping motif annotations, we collapsed similar overlapping motifs into a single interval.We defined motif similarity based on membership within the same matrix cluster, as designated by JASPAR 2024 (Rauluseviciute et al. 2023) (Table S1). The 377 motifs with conserved UMR enrichment represented 35 unique clusters. We manually annotated each of the 35 clusters with a descriptor of the motifs contained within (e.g. “HSF/GARP” if the cluster contained motifs from HSF and GARP TFs.) After collapsing overlapping motif instances by cluster, we quantified the number of cluster instances per upstream region for each assembly.

*Cis-regulatory feature overlap*: We quantified overlap of scATAC-seq peaks, MOA-seq peaks, and conserved non-coding sequences with 278,610 motif instances spanning 500 bp upstream regions in maize B73 v5. Merged scATAC peaks for all maize cell types were downloaded from

https://github.com/plantformatics/maize_single_cell_cis_regulatory_atlas/raw/refs/heads/master/ all_ACRs.celltype_calls.overlap.bed.gz (Marand et al. 2021) and uplifted to maize B73 v5 coordinates using CrossMap v.0.7.3 (Zhao et al. 2014) and a chain file downloaded from MaizeGDB. 168,009 peaks were successfully lifted over and used for the overlap analysis.

Additionally, 701,701 merged MOA-seq peaks for 21 maize hybrids projected onto B73 coordinates were obtained from (Engelhorn et al. 2023) and 1,664,343 conserved non-coding sequences for the Andropogoneae clade were obtained from (Stitzer et al. 2025). We used the intersect function from bedtools v2.29.2 (Quinlan and Hall 2010) with -u to quantify how many motifs overlapped each feature type. Null motif overlap values were calculated using 100 shuffled intervals for each feature type, using bedtools shuffle with -noOverlapping to prevent shuffled intervals from overlapping

*Motif abundance and variability:* We used our motif counts in 500 bp upstream regions to estimate 1) the median occurrence rate and 2) coefficient of variation of each motif cluster across assemblies. We calculated occurrence rates of each motif for each assembly by dividing the total number of instances of a particular motif in 500 bp upstream regions by the number of 500 bp upstream regions represented in that assembly, effectively calculating the average number of motif instances per upstream region. We then calculated the median and coefficient of variation for motif occurrence rates across all assemblies.

*Conservation of motif instances:* Motif conservation was defined for each assembly relative to Maize B73, with the percent conservation of maize motifs at a single orthogroup defined as 1 - (# motifs present at the target assembly gene / # of motifs present at the maize ortholog). As with most estimates of TFBS turnover, we did not consider motifs present in other lineages but absent in maize. For example, conservation at a single ortholog with three motif types present might be quantified as follows:

|  | Motif A instances | Motif B instances | Motif C instances | Total maize motifs present |
| --- | --- | --- | --- | --- |
| Target assembly | 1 | 0 | 2 | 2 |
| Maize | 1 | 1 | 1 | 3 |
| % conservation | 100% | 0% | 100% | 67% |

We calculated motif conservation relative to maize both at individual orthogroups (“local”) and across all 12,784 orthogroups present in maize (“global”). For local and global conservation, we plotted motif conservation against genetic distance (calculated using the Angiosperms353 loci as described above). We fit exponential decay curves of the form

to the relationship between genetic distance (x) and motif conservation (y).

To identify functional enrichments for orthogroups with high and low motif conservation, we calculated the mean percent conservation across taxa for maize motifs. Using the mean conservation values for each orthogroup, we identified orthogroups within the top and bottom quartiles of motif conservation. The top and bottom quartiles corresponded to 3051 and 3031 maize genes, which we mapped to orthogroups using miniProt (v0.13.0) between the maize sequences and the reconstructed ancestral sequences of the orthogroups in this study. We then performed GO enrichment using the R/topGO package (Alexa and Rahnenführer 2009) with the “weight01” algorithm, which considers hierarchical relationships among terms. For background genes, we used the remaining ∼9700 orthogroups not contained in the target quartile. Using maize B73 v5 GO annotations downloaded from MaizeGDB (Woodhouse et al. 2021), we tested for significant cellular component, biological process, and molecular function terms and calculated p values from Fisher statistics.

To compare motif conservation at TF vs non-TF genes, we downloaded a list of maize TFs from Grassius (Gavgani et al. 2023) corresponding to 1,204 distinct orthogroups. Using orthogroup-level measures of motif conservation, we used an asymptotic two-sample Kolmogorov-Smirnov test to evaluate the difference in conservation between TFs and non-TFs (all orthogroups not contained in the Grassius list).

To compare motif conservation at genes with ChIP-seq support, we downloaded ChIP-seq data from four maize TFs (Tu et al. 2020) matched to motifs used in our analyses: ereb17 (MA1818.1), glk53 (MA1830.2), bhlh47 (MA1834.2), and hb34 (MA1824.2). For each TF, we identified a set of orthogroups for which the 500 bp upstream region of the maize ortholog intersected with a ChIP-seq peak. Then, for the motif clusters corresponding to each ChIP-seq TF we quantified conservation of motif instances between maize and every assembly across 1) all orthogroups 2) all orthogroups intersecting a maize ChIP-seq peak.

*Environmental characterization:* We followed an environmental characterization pipeline previously described by (Hsu et al. 2024) to characterize the environmental niches of the studied species. Briefly, for each species, we retrieved the geographical coordinates of occurrence from BIEN (Maitner et al. 2017)and GBIF databases and obtained various environmental features in which the species occurs. We were able to characterize the habitat environments for 706 out of 727 taxa and thus excluded taxa where we were unable to curate environmental data from subsequent analysis (Hsu 2025). The variation in habitat environments amongst the diverse species were summarized into environmental principal components (“envPCs”). Each envPC captures a different combination of spatial and climatic patterns, with the multivariate decomposition process resulting in synthetic variables (the envPCs) that preserve important statistical properties, such as orthogonality, lack of autocorrelation, and normality. The top 10 environmental PCs were modeled as predictor variables in our phylogenetic mixed models.

*Cross-species environmental association models:* For models associating global motif occurrence rates with environment, we fit 35 phylogenetic mixed models of the following form with ASReml-R v4.2 (Butler et al. 2009), with one model per motif and one observation per assembly in each model:

The phylogenetic relationship matrix (phyloK) was fit as a random effect to control for shared evolution across species. All other predictor variables were fit as fixed effects. Occurrence rates for each focal motif were calculated as described in the “Motif abundance and variability*”* section above. We quantified global motif density (across all motifs) for each assembly by summing all 35 individual motif occurrence rates. Mono/di-nucleotide principle coordinates were used as covariates to control for background genome characteristics such as GC content, which influence motif detection rates. To estimate mononucleotide and dinucleotide frequencies in neutral genomic background regions, we extracted sequences for all introns shorter than 150bp using the miniProt alignments previously described in the “Orthogroup construction” section. We used the fasta-get-markov function from the MEME suite with -m 1 to calculate mono/di-nucleotide frequencies, and five principle coordinate axes were generated from these frequencies using the R/ade4 package (Dray and Dufour 2007). We fit the envPCs terms as fixed effects, along with the mono/di-nucleotide content and global motif density covariate terms. We calculated p values for each envPC term using Wald tests. We plotted the distribution of the 350 observed p values against p values expected under a uniform distribution to identify deviations from the null hypothesis.

For orthogroup-specific models of motif-environment associations, we used ASReml-R v4.2 to run phylogenetic mixed models of the following form, with one observation per assembly:

Environmental PCs (envPCs) were specified as fixed effects with the phylogenetic relationship matrix (phyloK) as a random effect to control for shared evolution across species. In total 498,287 models were run (# of motifs x # of orthogroups). Models with over 95% variance explained by phyloK were discarded to minimize unstable fixed effect estimates. Wald tests were performed on each envPC term to calculate p values. We plotted the distribution of the ∼ 5 million observed p values against p values expected under a uniform distribution to identify deviations from the null hypothesis of no association between environment and motif occurrence. To visualize how motif gain/loss is associated with temperature adaptation events, we reconstructed ancestral temperature values across the Poaceae phylogeny using the contMap function from phytools v2.3-0 (Revell 2024).

*Visual comparison of orthologous regions:* To compare orthologous upstream regions, we performed pairwise whole-genome alignments between *O. sativa* var. ZhenShan97 and *Zizania palustris* using Anchorwave v.1.2.2 (Song et al. 2022) with parameters -R 2 -Q 1. We generated chain files from Anchorwave alignments using a custom “MAFtoChain” script, then lifted over motifs into rice coordinates using CrossMap v0.7.3 (Zhao et al. 2014). To visualize open chromatin regions, we used RiceENCODE to download FAIRE-seq and RNA Polymerase II ChIP-seq tracks from (Zhao et al. 2020).

DATA AVAILABILITY: Short read genome assemblies will be made publicly available on Ag Data Commons upon publication. Code to generate the analyses and figures can be found at https://github.com/maize-genetics/poaceae_tfbs.

## ACKNOWLEDGEMENTS

<u>Funding:</u> This work was supported by NSF PanAnd Grant Award #1822330 and USDA-ARS Project Number 8062-21000-052-004-A. E.S.B. is supported by the USDA-ARS (ARS project number 8062-21000-052-000-D). A.J.S. was supported by NSF GRFP DGE – 2139899. M.C.S. was supported by NSF PRFB 1907343. A.P.M. is supported by NIH NIGMS 1R00GM144742. Computational resources were provided by the SCINet project and the AI Center of Excellence of the USDA Agricultural Research Service, ARS project numbers 0201-88888-003-000D and 0201-88888-002-000D. Additional computational resources and data management were provided by the Bioinformatics Facility (RRID:SCR_021757) at the Cornell Institute of Biotechnology.

<u>Collaborators:</u> We thank Julia Englehorn for early access to MOA-seq data. Thanks to Chelsea Specht, Charles Danko, Jian Hua, and the entirety of the Buckler lab for valuable feedback throughout the project.

## Germplasm collection

Obtaining plant material for this project would not be possible without our many collaborators in the field and herbaria. We’re grateful for their expertise, willingness to aid in field collecting, and sharing of precious herbarium material. In no particular order, we thank the following individuals, governmental organizations, and research institutions:

### Individuals

Taylor AuBuchon-Elder led the germplasm collection and plant maintenance with help from the following individuals: Donald Danforth Plant Science Center Plant Growth Facility staff, Rémy Pasquet, Cassiano Welker, Chrissy McAllister, Pat Minx, Bess Bookout, Maria Vorontsova, Jordan Teisher, Pete Lowry, Sarah Mathews, Richard Jobson, Tim Teetaert, Julie Pelc, Rev.

Dennis Testerman, Russel Juelg, James Cole, Ron Day, Courtney Angelo, Chris Matson, Douglas Rogers, Michael McKain, Marshall Shaw, Arthur Stiles, Chuck Byrd, Kyle Dillard, Robert Findling, Nancy Sferra, Jonathan Bailey, Lynn Riedel, Brian Anacker, Kirsti Harms, Brandon Crawford, Charlotte Reemts, Matt McCaw, Kevin Thuesen, Michelle Bertelsen, Ryan Middleton, Wesley Newman Plant vouchers from available material were deposited at Missouri Botanical Garden Herbarium; and for Australian wild collections, duplicates were deposited at Australia’s National Herbarium in Canberra.

### Herbaria (silica dried leaf tissue and herbarium specimen tissue)

Missouri Botanical Gardens, Royal Botanic Garden Kew, Muséum national d’Histoire naturelle, Australia National Herbarium, Queensland Herbarium, Northern Territory Herbarium, National Herbarium of New South Wales *Governmental organizations (permits, permissions, and live plant or seed material)* Australian National Government, New South Wales National Parks and Wildlife Service, Queensland Parks and Wildlife Service, Queensland Department of Environment Service, Northern Territory Government, Northern Territory Department of Natural Resources, Victoria Department of Environment, Land, Water, and Planning, North Carolina Department of Agriculture and Consumer Services, NC Plant Conservation Program, US National Park Service, US Department of Agriculture, City of Boulder Open Space and Mountain Parks, Florida Park Service and Department of Environmental Protection, City of Austin Water Quality Protection Lands

### Non-profits, NGOs, Research institutes (permits, permissions, and live plant material)

Katy Prairie Conservatory, Lady Bird Johnson Wildflower Center, The Nature Conservancy (Alabama, Arizona, Florida, Maine, Manitoba, Missouri, New Mexico, Texas), Native Prairie Association of Texas, University of Alabama, New Jersey Conservation Foundation, University of Puerto Rico at Mayagüez All collections were done in compliance with the Nagoya Protocol, with permits as required by local authorities.

We acknowledge and honor the many custodians and stewards of wild and domesticated grass diversity worldwide who have shaped the study, preservation, and cultivation of grasses.

