## Supplemental Appendix for "Extensive modulation of a conserved *cis*-regulatory code across 589 grass species"

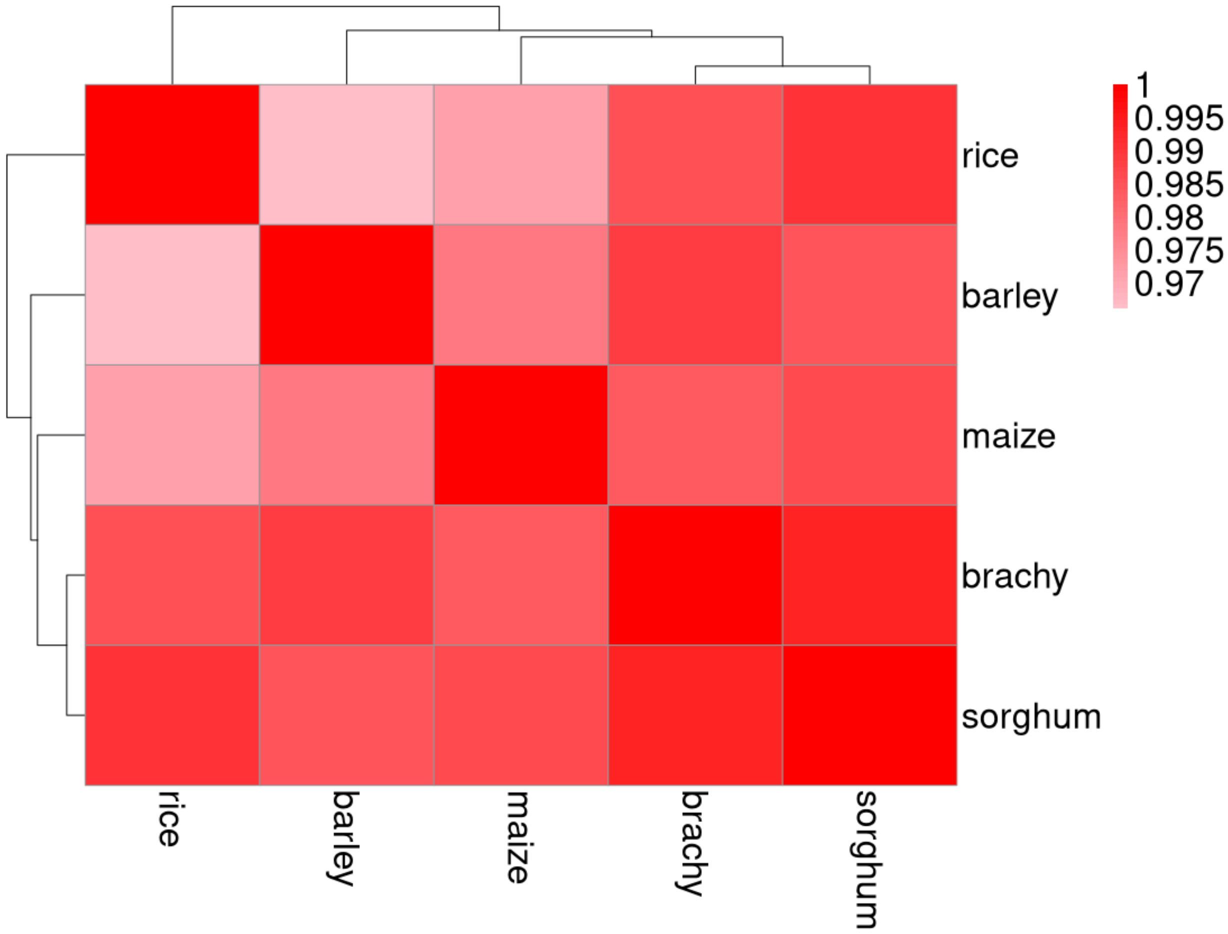


**Supplementary Figure 1. Fold-change UMR enrichment correlations across species.** Pearson correlations across 704 motifs are shown.

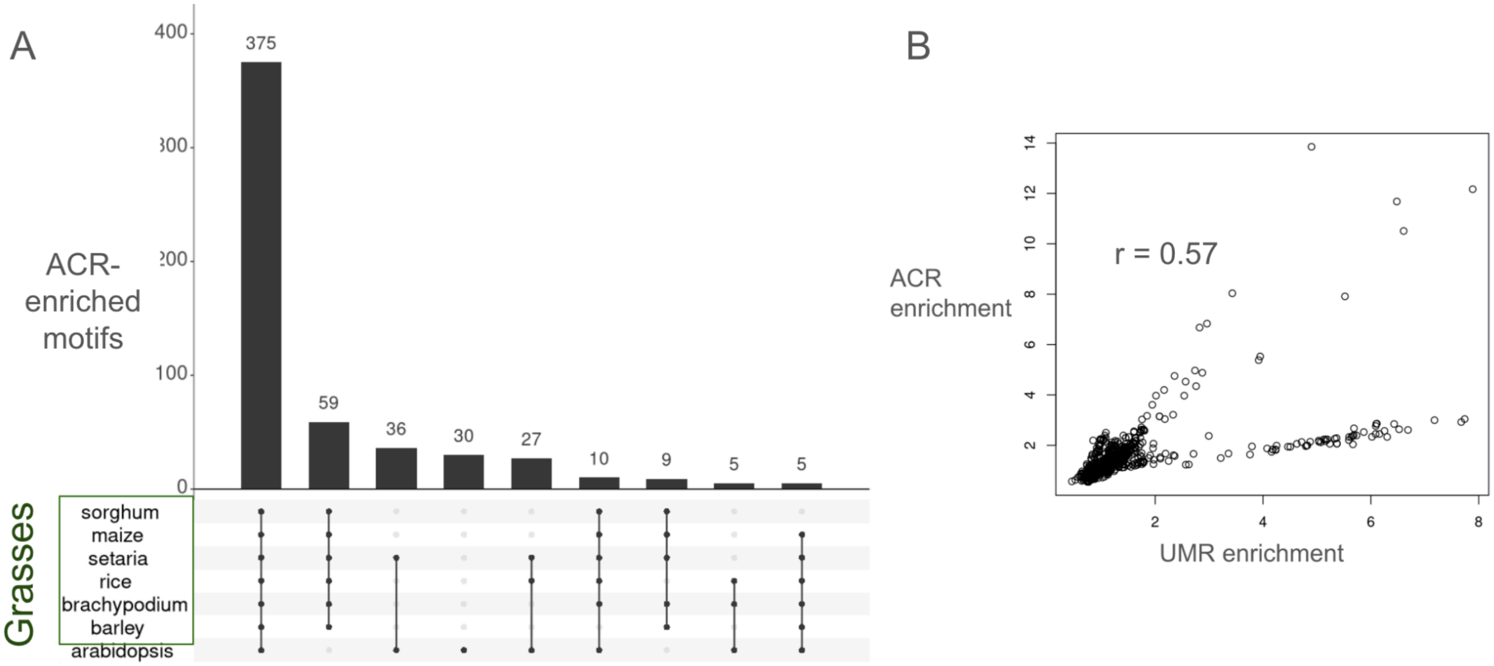


**Supplementary Figure 2. Motif enrichment in accessible chromatin regions.**

A) Enrichment of transcription factor motifs in unmethylated regions across species. The intersection bars show the number of enriched motifs for each species set. Intersections with fewer than five motifs are not shown. 336 / 377 (89%) of the shared UMR-enriched motifs were also commonly enriched in ACRs across grass species.

B) Pearson correlation between UMR enrichment fold-change and ACR enrichment fold change across motifs in maize

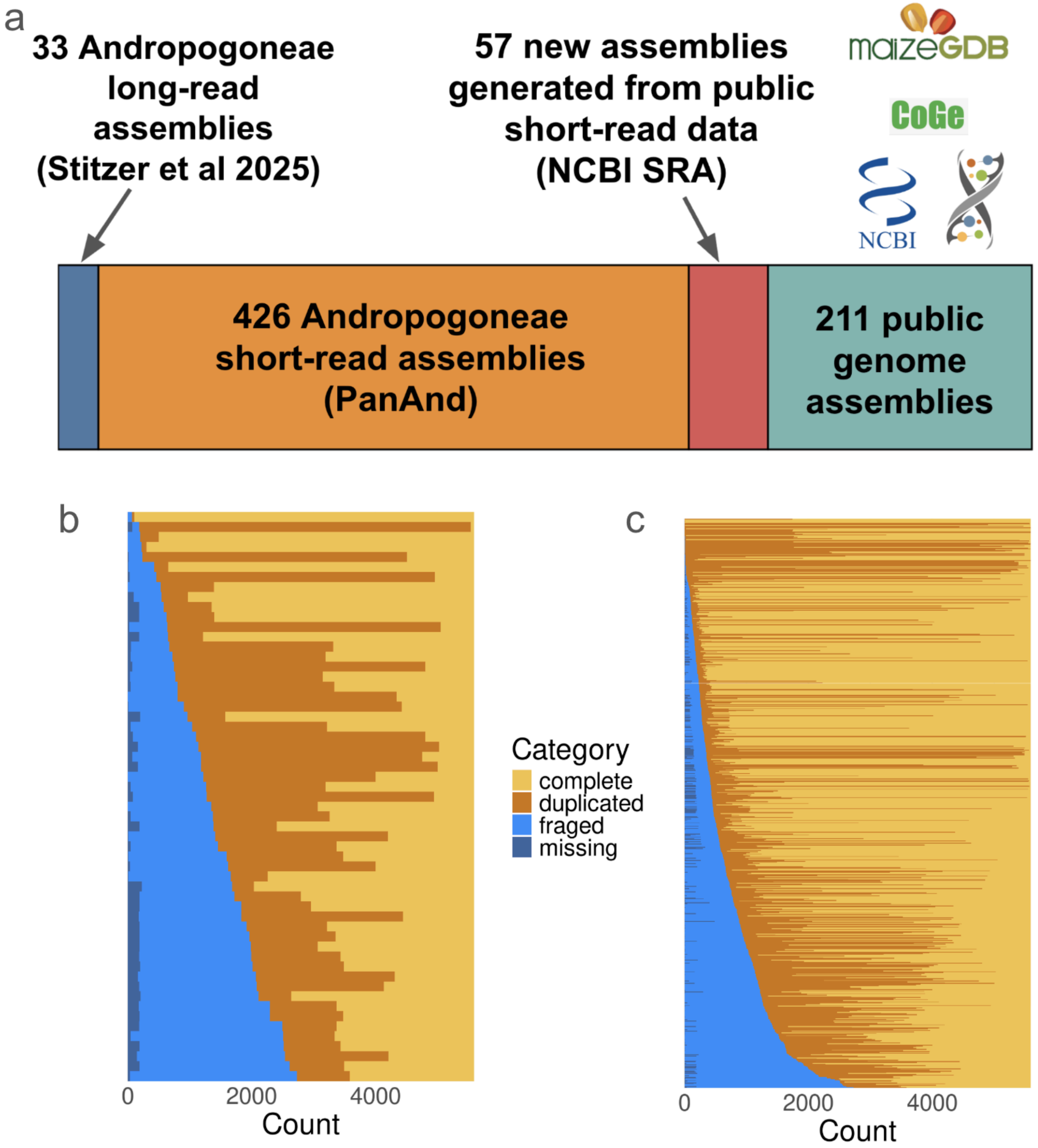


**Supplementary Figure 3. Breakdown of genome assemblies used in this study.**

a) Origins of 727 genome assemblies, representing 589 distinct species, that were used for analyses.
b) TABASCO scores representing assembly completeness across the 57 genome assemblies generated for this study from public SRA WGS reads.
c) TABASCO scores across all 727 assemblies.

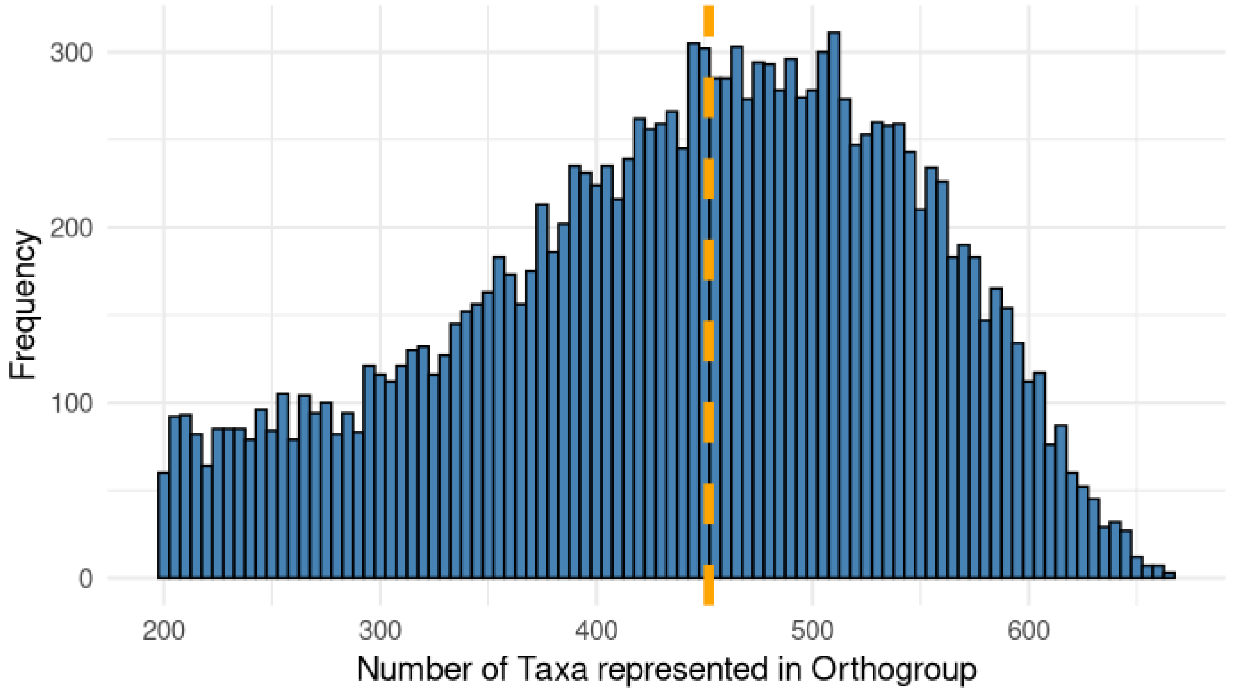


**Supplementary Figure 4. Taxa representation across orthogroup.** Dashed orange line denotes the median number of taxa represented across 15,555 orthogroups.

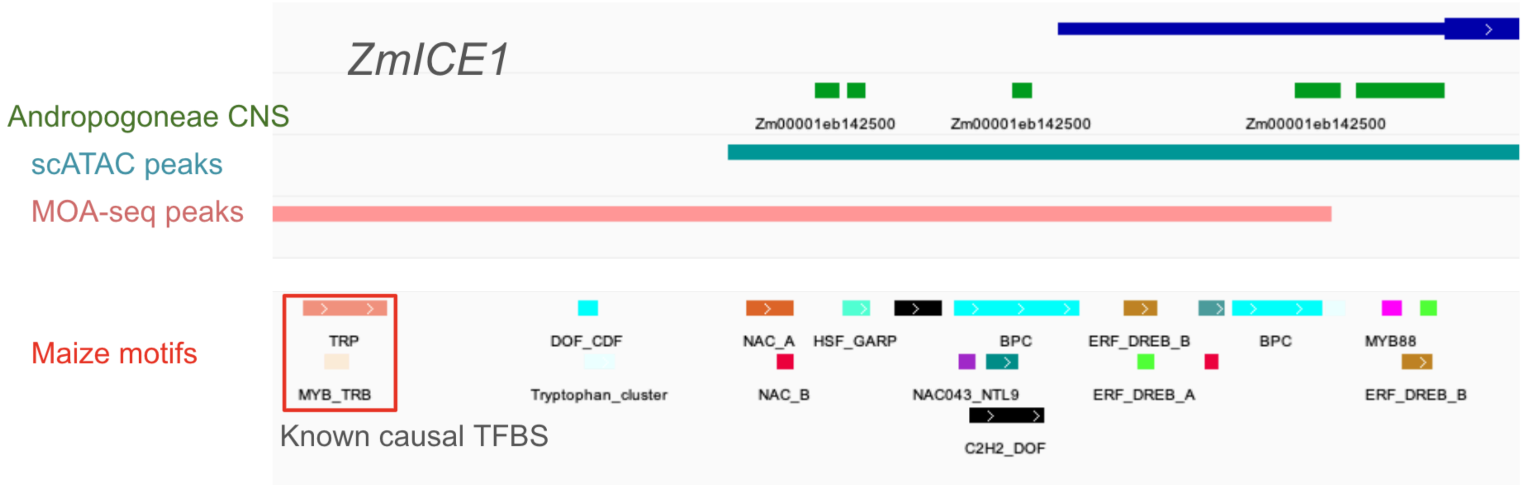


**Supplementary Figure 5. Example motif annotation upstream of *ZmICE1*.** A known causal TFBS variant from (Jiang et al. 2022) is highlighted in the red box.

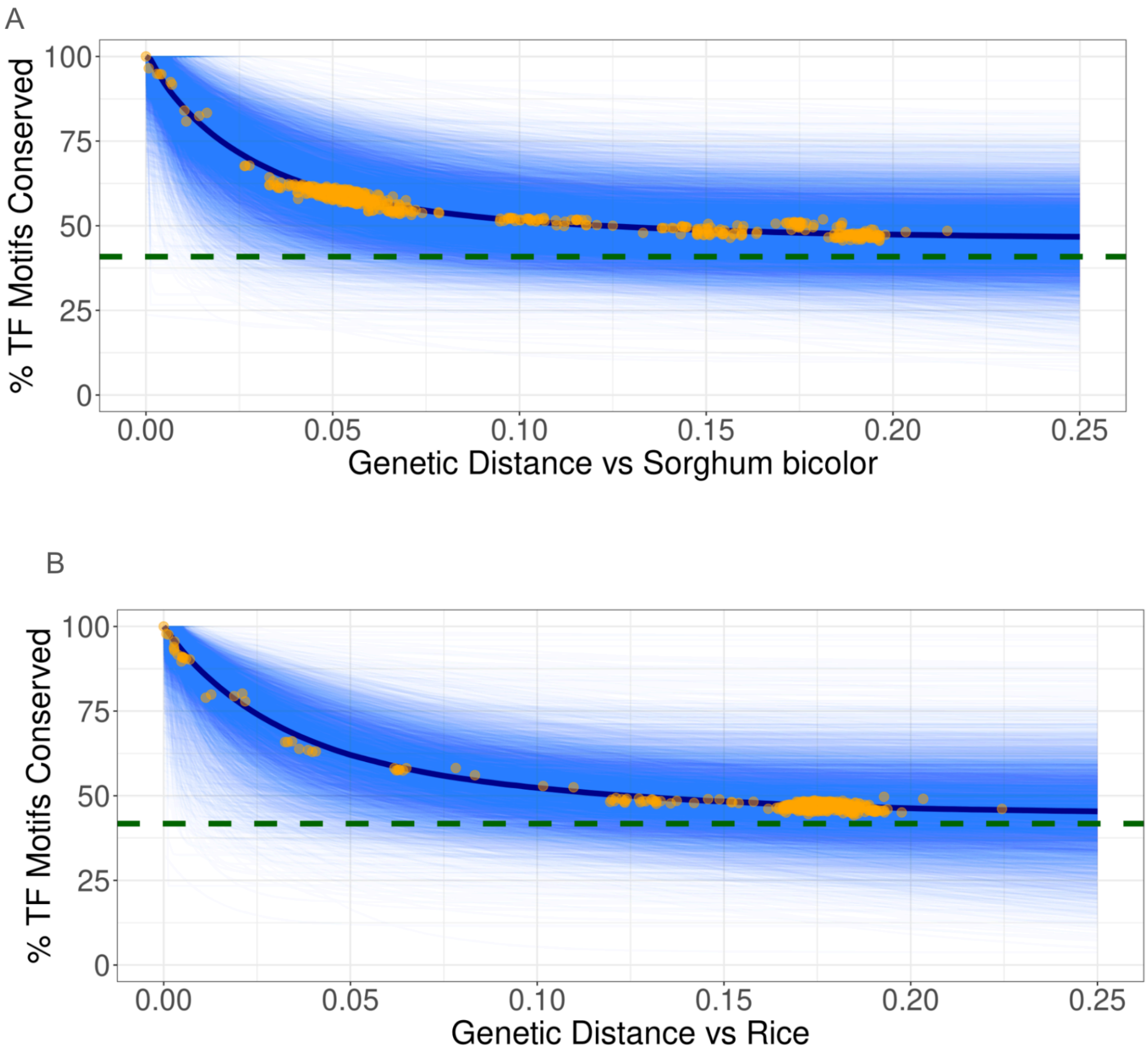


**Supplementary Figure 6. Decay in motif conservation relative to *Sorghum bicolor* (a) and *Oryza sativa* (b).** Genetic distance was estimated using pairwise distances between the focal species and the other Poaceae taxa at the Angiosperms353 loci. Blue lines represent exponential decay curves fit for each orthogroup. Orange points show the mean percentage of focal species motifs conserved in each Poaceae species across all orthogroups, with an exponential decay curve depicted in dark blue. Dashed green line represents the mean percentage of motifs conserved across 100,000 random pairs of genes in the focal species.

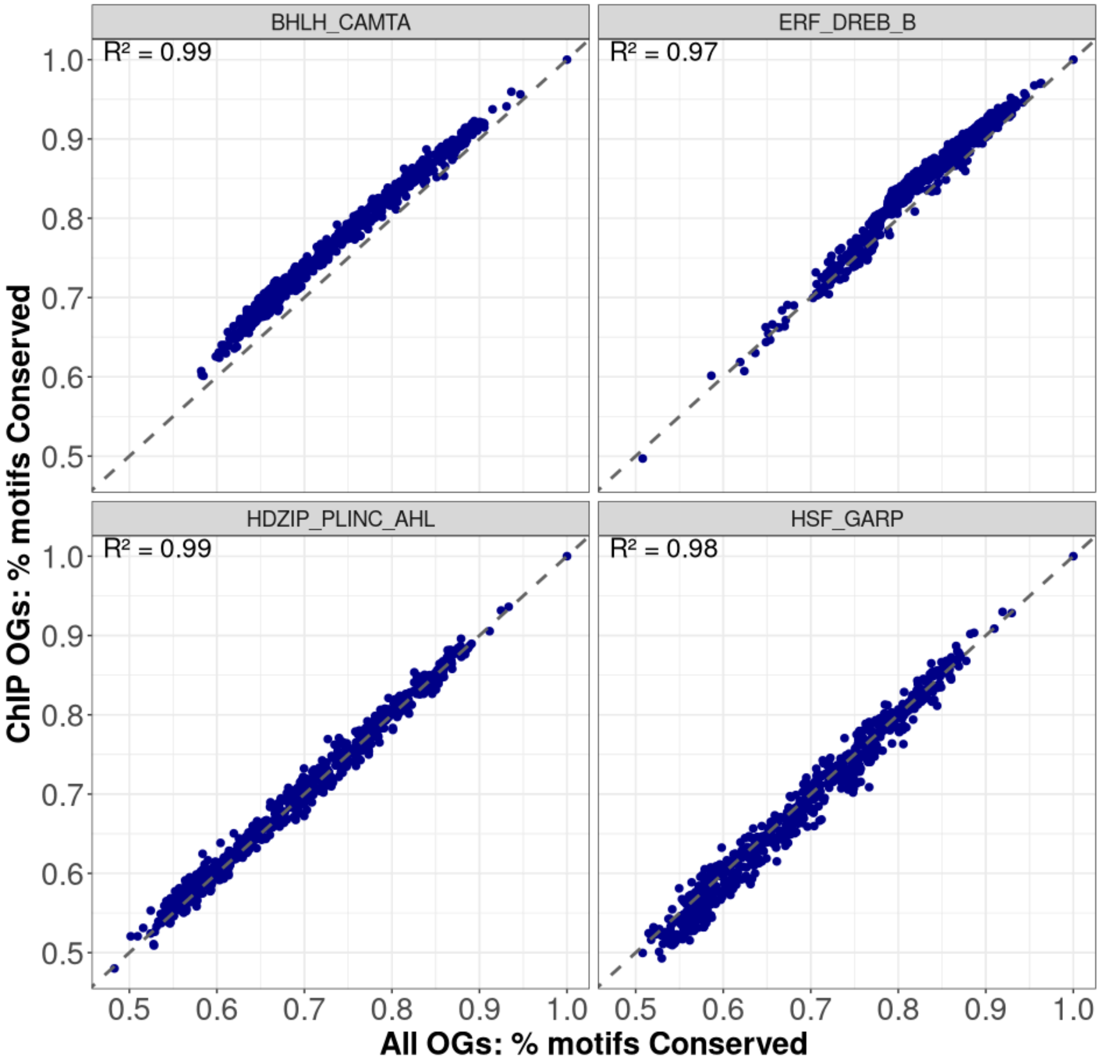


**Supplementary Figure 7. ChIP-seq comparison of motif conservation.** Motif conservation for four individual motif clusters (BHLH/CAMTA, ERF/DREB B, HDZIP/PLINC/AHL, HSF/GARP) is shown, plotting the mean conservation values across all orthogroups between maize and each of the other 726 taxa (x axis) with the mean conservation value across all orthogroups that intersected a matched ChIP-seq for the matched TF from maize (y axis). y=x line is shown in grey.

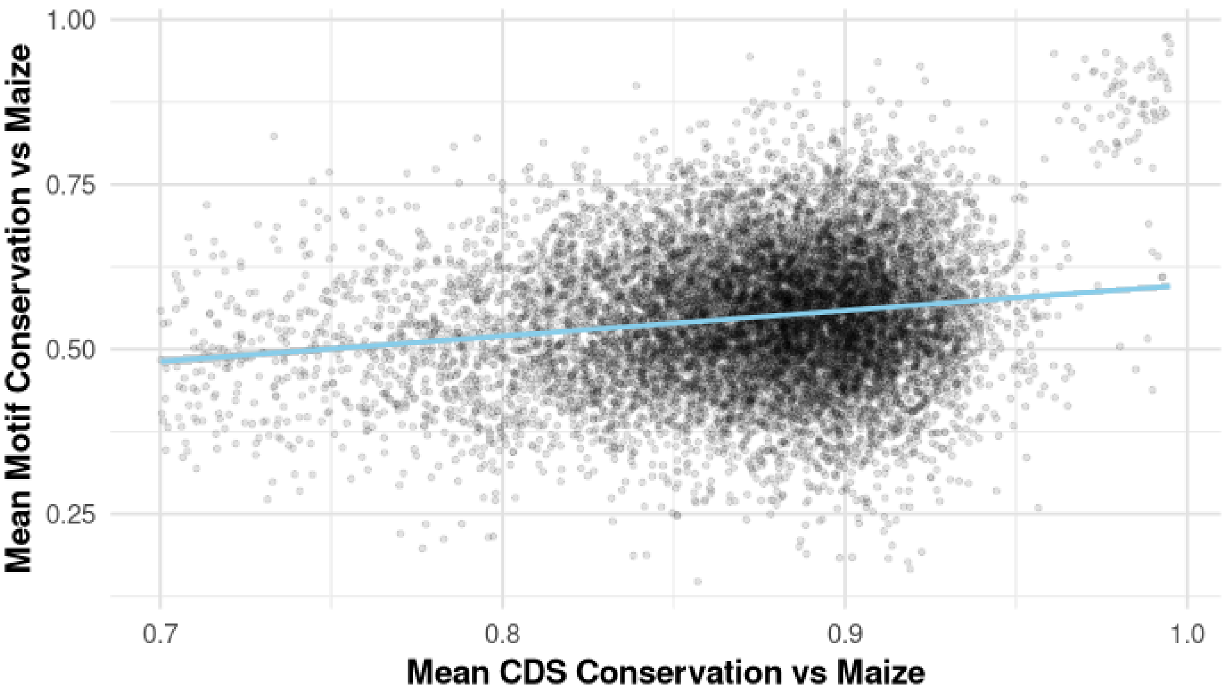


**Supplementary Figure 8. Motif vs coding sequence conservation across orthogroups.** For each orthogroup, the mean conservation values between maize and the 726 other taxa are depicted. A small number of orthogroups with <70% coding sequence conservation were excluded. The blue line depicts a linear model fit to the data with R-squared = 0.03.

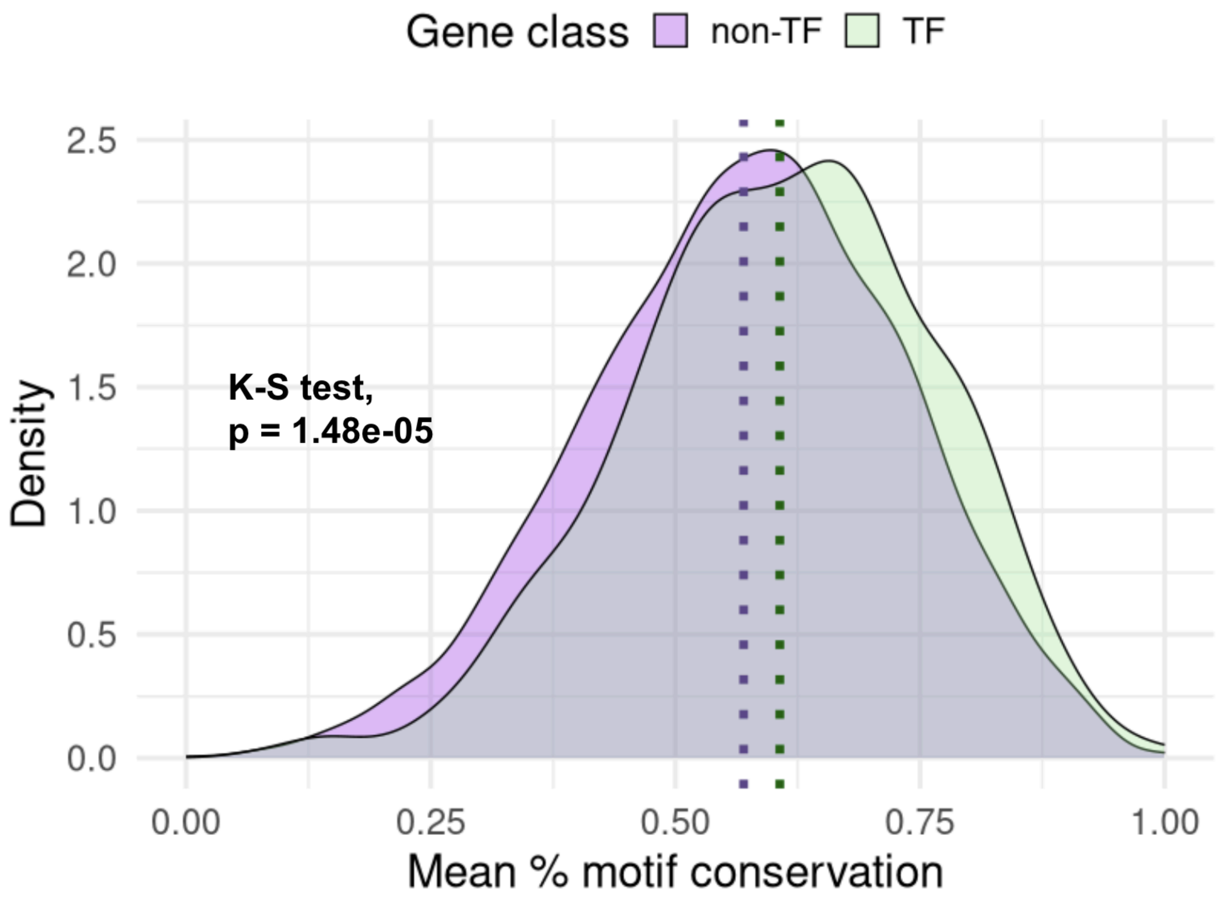


**Supplementary Figure 9: Maize vs sorghum motif conservation across syntenic orthologs**. Mean % motif conservation is shown calculated across 645 syntenic TF orthologs (“TF”) and 5414 background syntenic orthologs (“non-TF”).

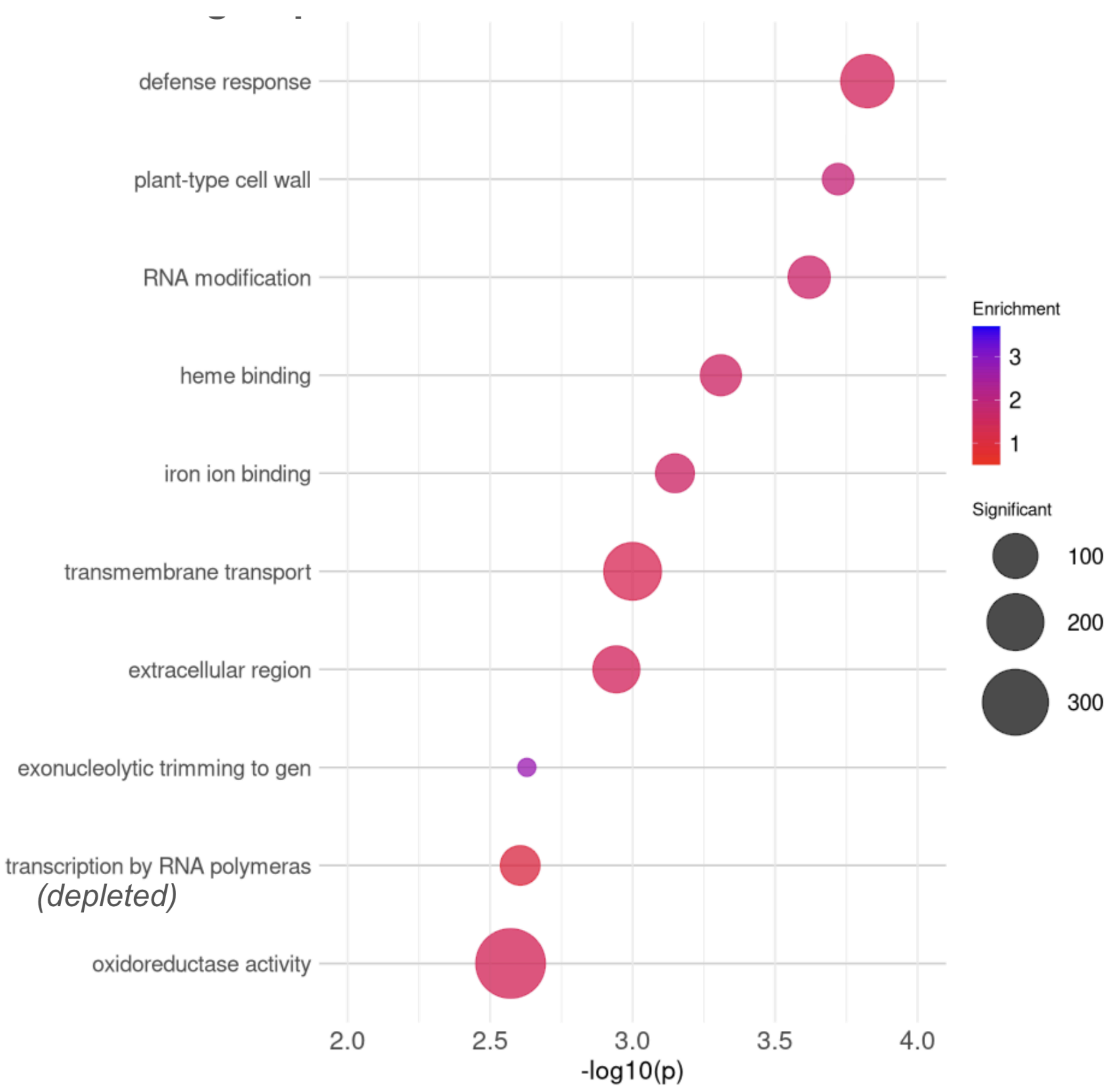


**Supplementary Figure 10: GO enrichment analysis for orthogroups with low motif conservation**. Enrichments were calculated for the orthogroups in the bottom quartile of motif conservation.
